# The protective effect of Capheic acid phenyl ester on hepatic ischemia-reperfusion injury in cholestatic rats

**DOI:** 10.1101/502294

**Authors:** Arif Aslaner, Tuğrul Çakır

## Abstract

**PURPOSE:** Caffeic acid phenyl ester (CAPE), which is an active component of propolis, has antioxidant, antiproliferative, immunomodulatory and anti-inflammatory properties and has been used for many years. The protective effect against ischemic reperfusion injury in the cholestatic liver is unknown and it is aimed to determine its effect in this study.

**MATERIALS AND METHODS:** Three groups of 18 Wistar albino rats were divided into cholestasis, control and study groups with bile duct attachment. Intraperitoneal administration of caffeic acid phenyl ester at the dose of 30 mg/kg/ day started 15 days ago and continued until the second operation in the study group. On the seventh day, hepatic ischemia was performed for 30 minutes followed by reperfusion for 60 minutes. Serum, plasma and liver samples were taken. Laboratory analysis, tissue glutathione, malondialdehyde, and myeloperoxidase levels were evaluated.

**RESULTS:** Significant reduction in the level of liver glutathione and a marked increase in malondialdehyde level and myeloperoxidase activity were observed in the cholestatic I / R group. After treatment, all parameters except serum bilirubin levels were reversed.

**CONCLUSIONS:** Intraperitoneal administration of CAPE may improve liver function and may reduce inflammation and oxidative stress in cholestatic I / R injury.

## 1. Introduction

Ischemia and reperfusion (I / R) injury for liver surgery is a common problem. After an ischemia, reperfusion causes the release of reactive oxygen radicals that are toxic to inducing lipid peroxidation, mitochondrial damage and apoptosis. 1

In the presence of cholestasis, it is more susceptible to damage.2 When bile flow is at a standstill, rapid damage to sinusoidal endothelial cells, neutrophil accumulation and activation of Kuppfer cells are induced.2 Active neutrophils and Kuppfer cells are the source of ROS and oxidative damage.3

CAPE is an active component of honey bee propolis extracts and has been used for many years. It has anti-inflammatory, immunomodulatory, antiproliferative and antioxidant properties and has been shown to inhibit lipo-oxygenase activities as it suppresses lipid peroxidation.4,5 To date, no studies have reported on the effect of CAPE on cholestatic liver ischemia reperfusion injury. The aim of this study was to evaluate the protection of CAPE against liver damage.

## 2. Material and Methods

The experimental procedures of this study were reviewed by Akdeniz University, Local Committee for Animal Research and Ethics and conducted at Akdeniz University Laboratory of Experimental Animals (2014/09/15). After the power analysis, the number of animals was determined. In this study, 18 Wistar Albino male rats (220-2700 g; 6 months) were used. The rats were maintained with a constant temperature of 22 hours, a 12-hour light / dark cycle and 60% humidity and ad libitum available in food and water. In the Sham group, the abdomen was closed without surgical procedure. I / R injury and intraperitoneal (Falma Chemical Co. St. Louis, MO) treated (I / R + CAPE) group, I / R damage and cholestasis model in addition to 30 mg / kg / day CAPE treatment was administered. 15 days before the first operation I / R + CAPE group 30 mg / kg / day CAPE was applied and intraperitoneally continued for 7 days. Sham and cholestatic I / R groups received equal amounts of saline for 21 days. The rats were given postoperative analgesia with acetaminophen (Paracetamol; Sigma-Aldrich Chemistry, Steinheim, Germany) at a dose of 50 mg / kg / day by oral gavage. In the other two groups, partial liver ischemia, portal vein and hepatic artery were closed for 30 minutes and then reperfused for 60 minutes.

Blood and liver tissue samples taken to evaluate the effect of CAPE; total bilirubin (Tbil), direct bilirubin (Dbil), alanine aminotransferase (ALT), aspartate aminotransferase (AST), lactate dehydrogenase (LDH) and alkaline phosphatase (ALP) glutathione (GSH), malondialdehyde (MDA), myeloperoxidase (MPO) activities Interleukin-1alpha (IL-1α), interleukin-6 (IL-6) and tumor necrosis factor-alpha (TNF-α).

For statistical analysis, SPSS 19.0 (IBM SPSS Statistics for Windows, IBM Corporation, Armonk, NY, USA) software was used. Kruskal-Wallis test was used to compare the three groups and Mann-Whitney U test was used when statistical significance was found. p <0.05 was accepted as statistically significant.

## 3. Results

ALT, AST, LDH, ALP, GGT and TBIL, DBIL serum levels were significantly higher in the I / R and I / R + CAPE groups compared to the sham group. The results of MDA, GSH and MPO are shown in Table 1. Tissue MDA and MPO levels were higher in both the I / R and I / R + CAPE groups than the sham group. The difference between the difference, I / R + CAPE and sham groups was not significant for MDA level (p = 0.05), there was a significant difference between Sham, I / R and I / R and I / R + CAPE (p = 0.001). Tissue GSH levels were lower in both the I / R and I / R + CAFE groups compared to the sham group. When I / R and I / R + CAPE groups were compared, tissue MDA and MPO levels were significantly higher in the I / R group than in the I / R + CAPE group and the tissue GSH level was significantly lower. Plasma proinflammatory cytokines, TNF-α and IL-1β levels were significantly higher in both I / R and I / R + CAPE group than in sham group (p <0.02). (Table 2)

**Table 1:**
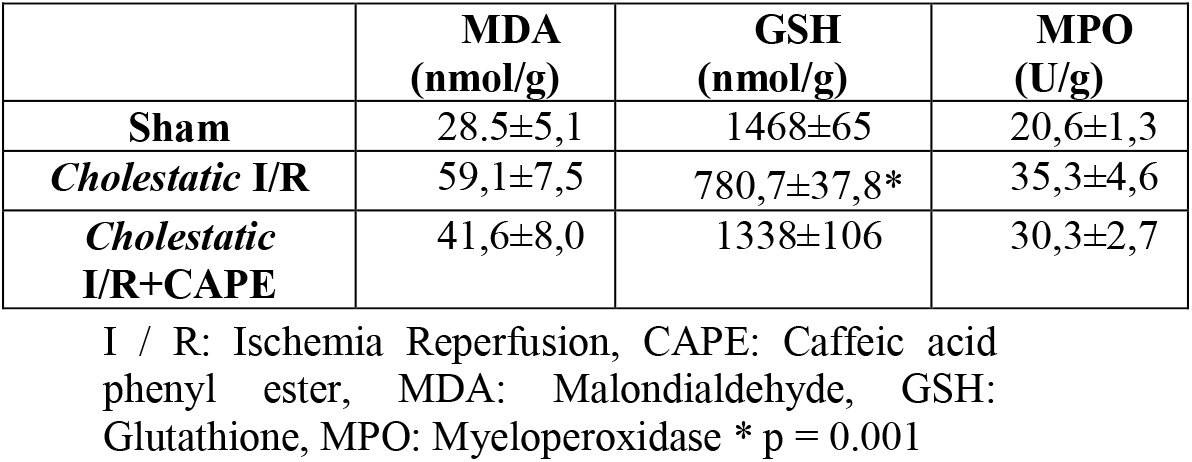
Tissue MDA, GSH and MPO levels of groups.

**Table 2:** The TNF-α and IL-1β levels of the groups.

| | TNF-α<br>(pg/ml) | IL-1β<br>(pg/ml) |
| --- | --- | --- |
| Sham | 9,9±1,5 | 208,5±26,8 |
| Cholestatic I/R | 22,3±4,7‡ | 410,0±37,7‡ |
| Cholestatic I/R+CAPE | 12,1±1,8 | 287,3±38,2 |

I / R: Ischemia Reperfusion, CAPE: Caffeic acid phenyl ester, TNF: Tumor Necrosis Factor, IL: Interleukin, ‡ $p < 0.02$

## 4. Discussion

The results of this study show that CAPE treatment significantly improves liver damage against cholestatic liver exposed to ischemia reperfusion injury. In the cholestatic I / R model, hepatocytes, sinusoidal endothelial cells and biliary ductal epithelium are damaged. The presence of these damages was demonstrated by a significant increase in AST, ALT, ALP and GGT in the cholestatic I / R group.

Ischemia causes metabolic changes in hepatocytes, especially in mitochondria. After reintroduction of oxygen to ischemic tissues, toxic reactive oxygen species (ROS) are released from mitochondria, mainly as the main causes of I / R injury.6

Cholestasis is another cause of oxidative stress damage. Blockage of biliary flow results in the accumulation and activation of polymorphonuclear neutrophils and Kuppfer cells that induce oxidative stress damage.3 The liver is more susceptible to I / R damage in the presence of obstructive jaundice.2 Therefore, an antioxidant such as CAPE is tested in a more harmonious state with the clinical application of the power of a drug We thought it should be.

CAPE, a flavonoid-like compound and an active ingredient in honey bee hives, propolis, has been used in traditional medicine for decades. CAPE is a flavonoid-like compound, anti-inflammatory, antioxidant, antiviral and anti-carcinogenic. Previous studies have demonstrated the properties of CAPE.8,9 Lipid peroxidation and inhibition of xanthine activity is via oxidase and nitric oxide synthase.

The glutathione form (GSH) is an essential component of the cellular defense mechanism against ROS-induced oxidative stress in rats with bile duct ligation. 12 Reduced GSH is oxidized by the enzymatic action of gluthation peroxidase, to remove ROS and to induce oxidation of glutathione in hepatocytes.6 CAPE application, GSH levels increased significantly and protective effects against oxidative damage were able to maintain glutathione levels. Less tissue GSH levels mean more oxidative stress damage.13 In our study, CAPE treatment provides a significant reduction of oxidative stress damage in the cholestatic I / R injury model.

MDA is a secondary product of oxidative stress during lipid peroxidation.14 In the current study, I / R damage caused a large amount of MDA accumulation in the liver. According to our results, tissue MDA levels were effectively reduced by CAPE.

The level of tissue MPO is directly proportional to the number of activated neutrophils and Kupffer cells used to assess liver I / R injury.15 Our results are consistent with previous studies of high MPO levels after icter. it may possibly be due to the antioxidant, free radical scavenging effect4,5 and inhibition of neutrophil infiltration.18

Activated Kupffer cells secrete TNFα and IL-1β cytokines. This is reflected in the plasma levels of TNFα and IL-1β, which is an indicator of both oxidative stress and inflammatory reaction.13 In the present study, cholestatic I / R damage caused a significant increase in plasma TNFα and IL-1β levels. This finding indicates that CAPE has a protective effect on cholestatic liver damage by inhibiting increased neutrophil infiltration into liver tissue.

In this study, a significant increase in proinflammatory cytokines in blood circulation and oxidative stress in ischemia reperfusion in icteric rats is observed. It has been shown that intraperitoneal administration of CAPE can improve liver function and reduce inflammation and oxidative stress in cholestatic I / R injury and is an effective protective agent against hepatic I / R damage in cholestatic liver. However, further studies are needed to evaluate the antioxidant, anti-inflammatory and liver protective effect of CAPE in clinical and experimental models.

## 5. Acknowledge

This study was financed by the Training and Research Planning Fund of Antalya Training and Research Hospital, with the permission of the Education and Research Planning Board at the same center.

